# Telomerase reverse transcriptase induces targetable alterations in glutathione and nucleotide biosynthesis in glioblastomas

**DOI:** 10.1101/2023.11.14.566937

**Authors:** Suresh Udutha, Céline Taglang, Georgios Batsios, Anne Marie Gillespie, Meryssa Tran, Sabrina M Ronen, Johanna ten Hoeve, Thomas G Graeber, Pavithra Viswanath

**Affiliations:** Department of Radiology and Biomedical Imaging, University of California San Francisco, San Francisco, CA, USA; Department of Molecular and Medical Pharmacology, University of California Los Angeles, Los Angeles, CA, USA; Crump Institute for Molecular Imaging; UCLA Metabolomics Center

**Keywords:** Telomerase reverse transcriptase, glutamine metabolism, glutathione, nucleotide biosynthesis, gliomas, metabolic synthetic lethality, cancer, metabolomics, *in vivo* stable isotope tracing

## Abstract

Telomerase reverse transcriptase (TERT) is essential for glioblastoma (GBM) proliferation. Delineating metabolic vulnerabilities induced by TERT can lead to novel GBM therapies. We previously showed that TERT upregulates glutathione (GSH) pool size in GBMs. Here, we show that TERT acts via the FOXO1 transcription factor to upregulate expression of the catalytic subunit of glutamate-cysteine ligase (GCLC), the rate-limiting enzyme of *de novo* GSH synthesis. Inhibiting GCLC using siRNA or buthionine sulfoximine (BSO) reduces synthesis of ^13^C-GSH from [U-^13^C]-glutamine and inhibits clonogenicity. However, GCLC inhibition does not induce cell death, an effect that is associated with elevated [U-^13^C]-glutamine metabolism to glutamate and pyrimidine nucleotide biosynthesis. Mechanistically, GCLC inhibition activates MYC and leads to compensatory upregulation of two key glutamine-utilizing enzymes i.e., glutaminase (GLS), which generates glutamate from glutamine, and CAD (carbamoyl-phosphate synthetase 2, aspartate transcarbamoylase, dihydroorotatase), the enzyme that converts glutamine to the pyrimidine nucleotide precursor dihydroorotate. We then examined the therapeutic potential of inhibiting GLS and CAD in combination with GCLC. 6-diazo-5-oxy-L-norleucin (DON) is a potent inhibitor of glutamine-utilizing enzymes including GLS and CAD. The combination of BSO and DON suppresses GSH and pyrimidine nucleotide biosynthesis and is synergistically lethal in GBM cells. Importantly, *in vivo* stable isotope tracing indicates that combined treatment with JHU-083 (a brain-penetrant prodrug of DON) and BSO abrogates synthesis of GSH and pyrimidine nucleotides from [U-^13^C]-glutamine and induces tumor shrinkage in mice bearing intracranial GBM xenografts. Collectively, our studies exploit a mechanistic understanding of TERT biology to identify synthetically lethal metabolic vulnerabilities in GBMs.

**SIGNIFICANCE:** Using *in vivo* stable isotope tracing, metabolomics, and loss-of-function studies, we demonstrate that TERT expression is associated with metabolic alterations that can be synergistically targeted for therapy in glioblastomas.

## INTRODUCTION

Glioblastoma (GBM) is the most common and lethal primary brain tumor in adults (1–3). Patient prognosis is bleak with a 5-year survival rate of 6.9% despite treatment with maximal safe surgical resection, radiation, and chemotherapy (1–3). Identifying targets that are intrinsically linked to GBM proliferation is a key challenge in the development of novel GBM therapies. Like most tumors, GBMs rely on telomerase reverse transcriptase (TERT) for telomere maintenance (4,5). Telomeres are cap-like nucleoprotein complexes that protect chromosomal ends from damage during replication (4,5). They shorten with every cell division, thereby imposing a natural limit on cell proliferation (4,5). TERT is the catalytic, rate-limiting component of the enzyme telomerase that synthesizes telomeric DNA (4,5). TERT expression is silenced in normal somatic cells, except stem cells, and reactivated in tumors cells via hotspot mutations in the *TERT* promoter that recruit the transcription factor GABP (5). Due to its essential role in proliferation, TERT is an attractive therapeutic target (6,7). However, silencing TERT delays growth but does not cause tumor regression in aggressive cancers like GBMs, partly due to the lag period before cell death due to telomere shortening occurs (6,7). Furthermore, while telomere uncapping agents such as 6-thio-2′-deoxyguanosine have shown promise in preclinical cancer models (8,9), drugs that directly inhibit TERT such as imtelstat have failed clinical trials due to toxicity towards stem and germline cells (6,7).

TERT expression has been linked to metabolic reprogramming in cancer (10). Specifically, we and others have demonstrated that silencing TERT depletes glutathione (reduced; GSH), NADPH and NADH in multiple cancer models, including GBMs, oligodendrogliomas, melanoma, hepatocellular carcinoma, and neuroblastoma (11–16). We then showed that TERT negatively regulates the forkhead box O1 (FOXO1) transcription factor, which is a master regulator of metabolism (15,17). FOXO1, in turn, inhibits expression of nicotinamide phosphoribosyl transferase (NAMPT), the rate-limiting enzyme for NAD+ biosynthesis, and of glyceraldehyde-3-phosphate dehydrogenase (GAPDH), the glycolytic enzyme that converts NAD+ to NADH (15). As a result, TERT elevates steady-state NAD+, NADH and the NADH/NAD+ ratio. In another study, we showed that TERT drives NADPH generation by upregulating glucose flux via the pentose phosphate pathway in oligodendrogliomas (13). However, the mechanism by which TERT upregulates GSH remains unknown.

GSH is the most abundant antioxidant within mammalian cells and exists predominantly in the reduced state in the cytosol (18,19). GSH plays an essential role in cell signaling and defense by functioning as an enzyme cofactor, modulating protein function via glutathionylation, and detoxifying reactive metabolites and xenobiotics (18,19). Importantly, GSH mitigates oxidative stress by scavenging reactive oxygen species (ROS) and is converted to oxidized glutathione (GSSG) in this process. Conversion back to GSH is mediated by glutathione reductase (18,19). GSH is a tripeptide of glutamate, cysteine, and glycine (*γ*-glutamylcysteinylglycine). Although cells can import GSH, extracellular GSH concentrations are ∼3 orders of magnitude lower than intracellular concentrations, with the result that intracellular GSH largely arises from *de novo* synthesis (20). The first step and rate-limiting step in GSH synthesis is the formation of *γ*-glutamylcysteine by γ-glutamylcysteine ligase (GCL), which is a heterodimer of a catalytic subunit (GCLC) and a regulatory subunit (GCLM). GSH synthetase then conjugates *γ*-glutamylcysteine with glycine to form GSH (20).

Metabolic reprogramming that is induced by TERT expression provides therapeutic targets that are potentially amenable to small molecule inhibition. Therefore, the goal of this study was to delineate the molecular mechanism by which TERT maintains GSH homeostasis and to determine whether this process can be exploited for GBM therapy. Our studies indicate that TERT acts via FOXO1 to upregulate GCLC expression and *de novo* GSH synthesis from [U-^13^C]-glutamine in GBM cells. Genetic or pharmacological inhibition of GCLC reduces ^13^C-GSH synthesis but causes compensatory upregulation of the glutamine utilizing enzymes glutaminase (GLS) and carbamoyl-phosphate synthetase 2-aspartate transcarbamoylase-dihydroorotatase (CAD) in a MYC-dependent manner. Importantly, combined inhibition of GCLC, GLS and CAD is synthetically lethal and induces tumor regression by abrogating GSH and nucleotide biosynthesis in preclinical GBM models *in vivo*.

## MATERIALS AND METHODS

### Patient-derived models

GBM6 and U251 cells were isolated from isocitrate dehydrogenase wild-type glioblastoma male patients as previously described (21,22). Cells were maintained in Dulbecco’s modified Eagle’s medium with 10% fetal calf serum (15,21,22). Cell lines were routinely tested for mycoplasma contamination, authenticated by fingerprinting, and assayed within 6 months. Patient biopsies were obtained from the UCSF Brain Tumor Center Biorepository in compliance with written informed consent policy (13–15). Biopsy use was approved by the Committee on Human Research at UCSF and research was approved by the Institutional Review Board at UCSF according to ethical guidelines established by the U.S. Common Rule.

### Silencing and overexpression studies

Two non-overlapping siGENOME or Accell siRNA sequences (Dharmacon) against human TERT (D-003547-02, D-003547-03), FOXO1 (D-003006-05, D-003006-06), GCLC (A-009212-15, A-009212-16), MYC (D-003282-14, D-003282-15) or non-targeting siRNA (D-001206-14) were used to silence gene expression. For exogenous expression of CA-FOXO1, TERT+ cells were transiently transfected with human CA-FOXO1 (pcDNA3 Flag-FKHR-AAA mutant; Addgene) (15). CA-FOXO1 expression was verified at 72 h by western blotting for the FLAG tag as described previously (15). To silence FOXO1 in TERT-cells, cells were transfected with siTERT siRNA for 24 h, washed and then transfected with siFOXO1 siRNA. FOXO1 silencing was verified at 72 h by quantifying FOXO1 transcription factor activity as described below.

### Activity assays

Telomerase activity was confirmed using the TRAPeze® RT kit (Sigma) (13–16). Glutathione reductase activity (Abcam, #ab83461), glutaminase activity (Abcam, #284547), and caspase activity (Abcam, #ab39401) were measured using kits according to manufacturer’s instructions. FOXO1 transcription factor activity was measured using a kit (Abcam, #ab207204). Briefly, the assay measures binding of active FOXO1 to a DNA sequence containing the FOXO1 binding site. FOXO1 is detected using a primary antibody that recognizes an epitope of FOXO1 accessible only when the protein is active and bound to its target DNA. GCL activity was measured as described previously (23). Cells or tissue were lysed in buffer containing 20 mM Tris-HCl, 1 mM EDTA, 250 mM sucrose, 20 mM sodium borate and 2 mM serine buffer and added to a reaction mixture (400 mM Tris-HCl, 40 mM adenosine triphosphate, 40 mM L-glutamic acid, 30 mM cysteine, 2 mM EDTA, 20 mM sodium borate, 2 mM serine, 40 mM MgCl_2_). After incubation at 37°C for 15 min, the enzymatic reaction was stopped by precipitation of proteins with 200 mM 5-sulfosalicylic acid. Following incubation with 2,3-naphthalenedicarboxyaldehyde (NDA) in the dark at room temperature for 30 min, the product (ℽ-glutamylcysteine-NDA) was detected by fluorimetry (472 nm excitation/528 nm emission). Data was compared to a standard curve and expressed as mmoles per mg protein. The carbamoyl phosphate synthetase II activity of the multifunctional CAD enzyme was measured as described previously (24,25). A two-step assay was used to measure carbamoyl phosphate synthesis by coupling the carbamoyl phosphate synthetase reaction to that of ornithine transcarbamoylase and quantifying the product i.e., citrulline. Specifically, cell lysate was added to a reaction mixture (50 mm HEPES, 100 mm KCl, 10 mm ATP, 20 mm MgCl_2_, 20 mm NaHCO_3_, 1 mm dithiothreitol, 5 mm ornithine, 0.2 units of ornithine transcarbamoylase, and 10 mm glutamine, pH 7.6) and incubated at 37 °C for 20 min. After incubation, citrulline was quantified by colorimetry with diacetylmonoxime at 490 nm as previously described (25). Clonogenicity was measured using a soft agar assay according to standard procedures (26).

### Gene expression

Gene expression was measured by quantitative PCR and normalized to β-actin (27,28). The SYBR Green quantitative PCR kit (Sigma) was used with the following primers: *TERT* (forward primer: TCACGGAGACCACGTTTCAAA; reverse primer: TTCAAGTGCTGTCTGATTCCAAT), *FOXO1* (forward primer: GCAGCCAGGCATCTCATAA; reverse primer: CCTACCATAGCCATTGCAGC), *GCLC* (forward primer: ATGTGGACACCCGATGCAGTATT; reverse primer: TGTCTTGCTTGTAGTCAGGATGGTTT), *GCLM* (forward primer: GCCACCAGATTTGACTGCCTTT; reverse primer: CAGGGATGCTTTCTTGAAGAGCTT), *CAD* (forward primer: AGTGGTGTTTCAAACCGGCAT; reverse primer: CAGAGACCGAACTCATCCATTTC), *GLS* (forward primer: TGGACTATGAAAGTCTCCAACAAGA; reverse primer: CTCATTTGACTCAGGTGACACTTTT), *MYC* (forward primer: TCAAGAGGCGAACACACAAC; reverse primer: GGCCTTTTCATTGTTTTCCA), and β-actin (forward primer: AGAGCTACGAGCTGCCTGAC; reverse primer: AGCACTGTGTTGGCGTACAG).

### Metabolomics and stable isotope tracing in cells

∼5×10^5^ cells were seeded in 6-well plates and treated with vehicle (dimethyl sulfoxide, DMSO) or DON (5 μM), BSO (10 μM) or siRNA (20 μM) in regular cell culture media for 48 h (drug treatment) or 72 h (siRNA). For combination therapy, cells were treated with 1 μM each of DON and BSO for 72 h. For stable isotope tracing, cells were seeded and treated as above and incubated in media in which glutamine was replaced with [U-^13^C]-glutamine (99% enrichment; Cambridge Isotope Laboratories, final concentration 5.7 mM). Following incubation for 48 or 72 h, cells were washed with ice-cold ammonium acetate (150 mM, pH 7.3). 1 ml of pre-cooled methanol/water (80:20 v/v) was added to each well and plates incubated at –80 °C for 30 min. Cells were collected by scraping and centrifuged at 14,000 rpm for 15 min at 4 °C to remove debris. The supernatant was lyophilized, samples were reconstituted with 60 μL of pre-chilled acetonitrile/water (50:50, v/v), transferred into glass vials and utilized for liquid chromatography mass spectrometry (LC-MS) as described below.

### MRI

Animal studies were conducted in accordance with UCSF Institutional Animal Care and Use Committee guidelines. GBM6 cells were intracranially injected into female SCID mice using a stereotactic frame at 2 mm to the right of the medial suture, 2 mm behind the bregma and a depth of 2 mm. Tumor volume was determined by T2-weighted MRI using a preclinical 3T scanner and a T2 rapid acquisition with relaxation enhancement (RARE) sequence (TE/TR = 64/3700 ms, FOV = 30 x 30 mm^2^, matrix = 256 x 256, slice thickness = 1.5 mm, NA = 5). Once tumors reached a volume of 30 ± 5 mm^3^, this timepoint was considered day 0. Animals were randomized and treated with vehicle (saline) or a combination of JHU-083 (20 mg/kg) and BSO (20 mg/kg) daily via intraperitoneal injection. These doses were selected based on prior studies in preclinical cancer models (29,30). Mice were treated until they needed to be euthanized or the tumor was no longer visible on MRI. Animal survival was assessed by Kaplan-Meier analysis.

### 13C tracing *in vivo*

Stable isotope tracing experiments *in vivo* were performed using a previously established protocol (31) on mice bearing intracranial GBM6 tumors treated with vehicle (saline) or 20 mg/kg each of JHU-083 and BSO as described above. At day 7 post-treatment, mice were intravenously infused with [U-^13^C]-glutamine under anesthesia on a heating pad: a bolus of 428 mg/kg of [U-^13^C]-glutamine diluted in 0.18 ml in saline was injected within 1 min, and then, 14 mg/kg/min was infused for 2 h. At the end of the infusion, brain (tumor and contralateral) tissue was collected and snap-frozen. ∼15-25 mg of each tissue sample was homogenized in 1 ml of pre-cooled methanol/water (80:20 v/v) and centrifuged at 14,000 rpm for 10 min. The supernatant was then lyophilized and used for LC-MS as described below.

### LC-MS

LC-MS was performed using a Vanquish Ultra High-performance LC system coupled to an Orbitrap ID-X Tribrid mass spectrometer (Thermo Fisher), equipped with a heated electrospray ionization (H-ESI) source capable of both positive and negative modes simultaneously (32). Before analysis, the MS instrument was calibrated using calibration solution (FlexMix, Thermo Fisher). Cell or tissue samples along with blank controls were placed in the autosampler. Individual samples were run alongside a pooled sample made from an equal mixture of all individual samples to ensure chromatographic consistency. Chromatographic separation of metabolites was achieved by hydrophilic interaction liquid chromatography (HILIC) using a Luna 3 NH2 column (150 mm x 2.1 mm, 3 μm, Phenomenex) in conjunction with a HILIC guard column (Phenomenex, 2.1 mm). The column temperature and flow rate were maintained at 27°C and 0.2 mL/min respectively. Mobile phases consisted of A (5 mM ammonium acetate, 48.5 mM ammonium hydroxide pH 9.9) and B (100% Acetonitrile). The following linear gradient was applied: 0.0-0.1 min: 85-80% B, 0.1-17.0 min: 80-5% B, 17.0-24.0 min: 5% B, 24.0-25.0 min: 5-85% B, 25.0-36.0 min: 85% B. The injection volume and auto sampler temperature were kept at 5 μL and 4°C respectively. High-resolution MS was acquired using a full scan method alternating between positive and negative polarities (spray voltages: +3800kV/-3100kV; sheath gas flow: 45 arbitrary units: auxiliary gas flow: 15 arbitrary units; sweep gas flow: 1 arbitrary unit; ion transfer tube temperature: 275°C; vaporizer temperature: 300°C). Three mass scan events were set for the duration of the 36-minute run time. The first was the negative polarity mass scan settings at full-scan-range; 70-975 m/z. Positive polarity mass scan settings were split to two scan events, full-scan-range; 70-360 m/z and 360-1500m/z, in that order. MS1 data were acquired at resolution of 60,000 with a standard automatic gain control and a maximum injection time of 100 ms. Data was acquired using Xcalibur software. Chromatograms were reviewed using FreeStyle (Thermo Fisher) and a 5 ppm mass tolerance. Peak areas were quantified using TraceFinder from either positive or negative modes depending on previously run standards. Peak areas were corrected to blank samples and normalized to cell number or wet weight of tissue and the total ion count of that sample. For ^13^C isotope tracing, % enrichment for each isotopomer was calculated after correcting for natural abundance using Escher Trace (33).

### Statistical analysis

All experiments were performed on a minimum of 3 samples (n≥3) and results presented as mean ± standard deviation. Statistical significance was assessed in GraphPad Prism 10 using a two-way ANOVA or two-tailed Welch’s t-test with p<0.05 considered significant. Analyses were corrected for multiple comparisons using Tukey’s method, wherever applicable. *=p<0.05, **=p<0.01, ***=p<0.001 and ****=p<0.0001.

### Data availability

Data included in this manuscript that support the findings and conclusions of this study are available from the corresponding author upon reasonable request.

## RESULTS

### TERT drives GSH synthesis from glutamine by upregulating GCLC expression in GBMs

First, we examined the effect of silencing TERT using two non-overlapping siRNA sequences in GBM6 and U251 cells and compared to cells transfected with non-targeting control siRNA (Fig. 1A and Supplementary Fig. S1A). Consistent with our previous studies (13–15), GSH levels were significantly reduced in TERT-cells relative to TERT+ (Fig. 1B). Elevated steady-state GSH can result from higher synthesis or elevated recycling of GSSG to GSH. We confirmed that neither GSSG levels (Fig. 1C) nor glutathione reductase (Supplementary Fig. S1B) activity were altered in TERT-cells relative to TERT+. Since glutamine is a key carbon donor for *de novo* GSH synthesis (20,34), we then traced [U-^13^C]-glutamine metabolism in our models. [U-^13^C]-glutamine (m+5) is converted by glutaminase (GLS) to glutamate (m+5), which is conjugated with cysteine by GCL to produce ℽ-glutamylcysteine. The addition of glycine to ℽ-glutamylcysteine produces GSH (m+5). As shown in Fig. 1D, TERT silencing significantly reduced ^13^C labeling of GSH in both GBM6 and U251 models. Concomitantly, GCLC expression and GCL activity were significantly reduced in TERT-cells relative to TERT+ (" ref-type="fig">Fig. 1E-1F). We did not observe any change in expression of GCLM or glutathione synthetase in TERT-cells relative to TERT+ (Supplementary Fig. S1C-S1D).

**Figure 1.**
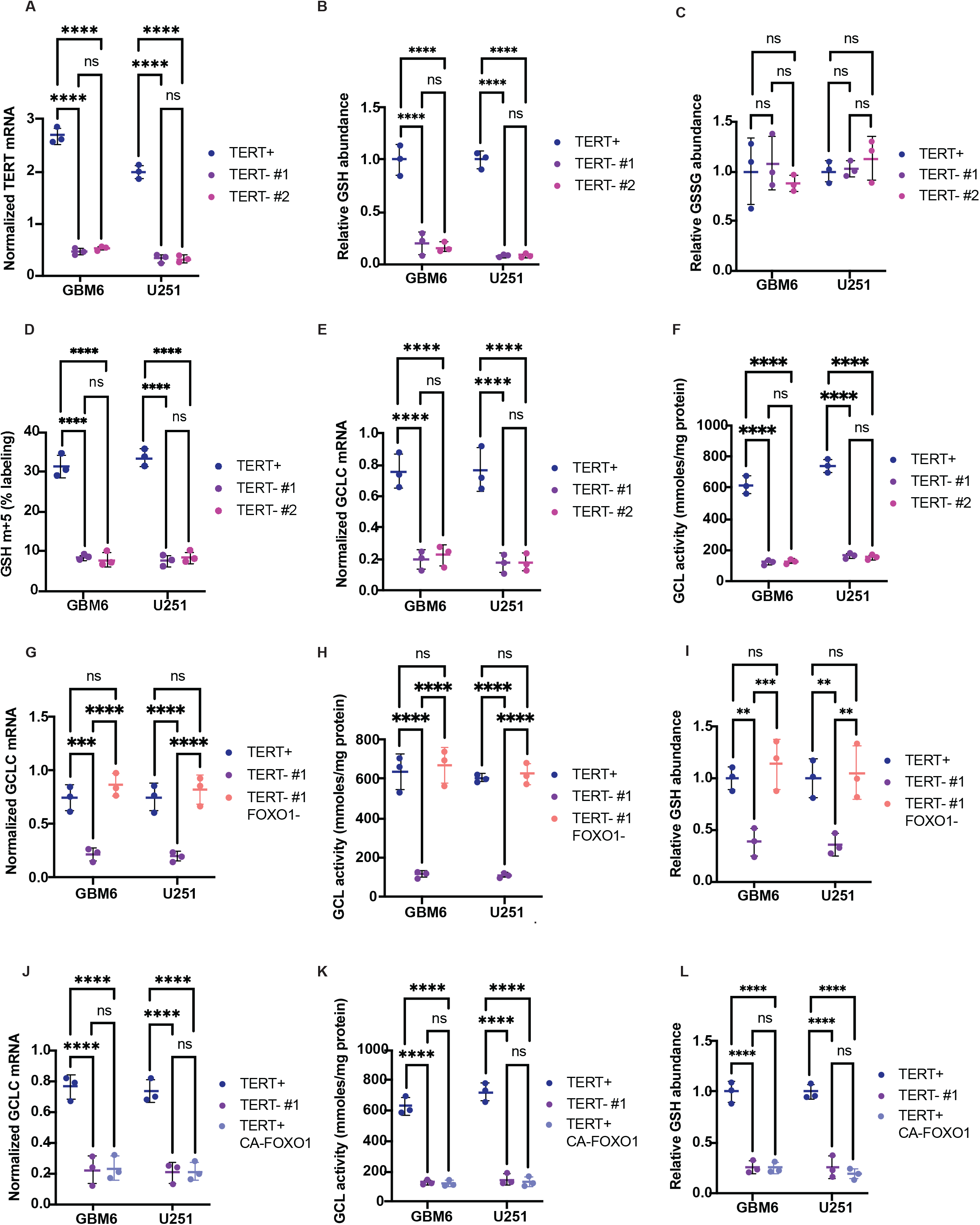
TERT acts via the FOXO1 transcription factor to upregulate GCLC, the rate-limiting enzyme in GSH synthesis, in GBM cells. TERT mRNA (A), GSH pool size (B), GSSG pool size (C), % ^13^C labeling of GSH from [U-^13^C]-glutamine (D), GCLC mRNA (E), and GCL activity (F) in GBM6 and U251 cells transfected with siRNA against a non-targeting control (TERT+) or two non-overlapping sequences against TERT (TERT-#1 and TERT-#2). Effect of silencing FOXO1 in TERT-cells on GCLC mRNA (G), GCL activity (H), and GSH pool size (I) in GBM6 and U251 cells. Effect of expressing a constitutively active form of FOXO1 (CA-FOXO1) in TERT+ cells on GCLC mRNA (J), GCL activity (K), and GSH pool size (L) in GBM6 and U251 cells. ** represents p<0.01, *** represents p<0.001 and **** represents p<0.0001.

Next, we examined the mechanism by which TERT upregulates GCLC expression in GBMs. The forkhead box O (FOXO) family of transcription factors coordinate metabolism and redox homeostasis (35), and we previously showed that TERT negatively regulates FOXO1 in GBMs via inhibitory phosphorylation (15). Phosphorylation sequesters FOXO1 in the cytoplasm, thereby preventing transactivation of FOXO1 target genes in the nucleus (35,36). To determine whether FOXO1 regulates GCLC expression, we examined the effect of silencing FOXO1 in TERT-cells (Supplementary Fig. S1E). As shown in " ref-type="fig">Fig. 1G-1I, silencing FOXO1 mimicked TERT expression and restored GCLC expression, GCL activity, and steady-state GSH to levels observed in TERT+ cells. Conversely, we examined the effect of expressing a constitutively active form of FOXO1 (hereafter named CA-FOXO1) in TERT+ cells (36). CA-FOXO1 is a FLAG-tagged form of FOXO1 that has been rendered constitutively active via mutation of all three phosphorylation sites to alanine (36). We confirmed that CA-FOXO1 restored FOXO1 transcription factor activity in TERT+ cells to levels observed in TERT-cells (Supplementary Fig. S1F). Expression of CA-FOXO1 in TERT+ cells mimicked TERT silencing and significantly reduced GCLC expression, GCL activity and GSH pool size in both GBM6 and U251 cells (" ref-type="fig">Fig. 1J-1L).

To validate the clinical relevance of our results, we examined GBM patient tissue samples and compared to non-neoplastic gliosis (since obtaining normal brain tissue is challenging) (13–15). We also examined tissue from patients with isocitrate dehydrogenase mutant astrocytoma, which use the alternative lengthening of telomeres pathway for telomere maintenance (14,37). We confirmed that TERT expression and telomerase activity were significantly higher while FOXO1 transcription factor activity was significantly lower in GBM biopsies relative to gliosis and astrocytoma (" ref-type="fig">Fig. 2A-2C), consistent with prior studies (15). As shown in " ref-type="fig">Fig. 2D-2F, GCLC expression and GCL activity were significantly higher in GBM patient tissue relative to gliosis or astrocytoma, and TERT expression was significantly correlated with GCLC expression in GBM biopsies. Collectively, our results suggest that TERT relieves FOXO1-mediated repression of GCLC expression, thereby upregulating GSH synthesis in GBMs.

**Figure 2.**
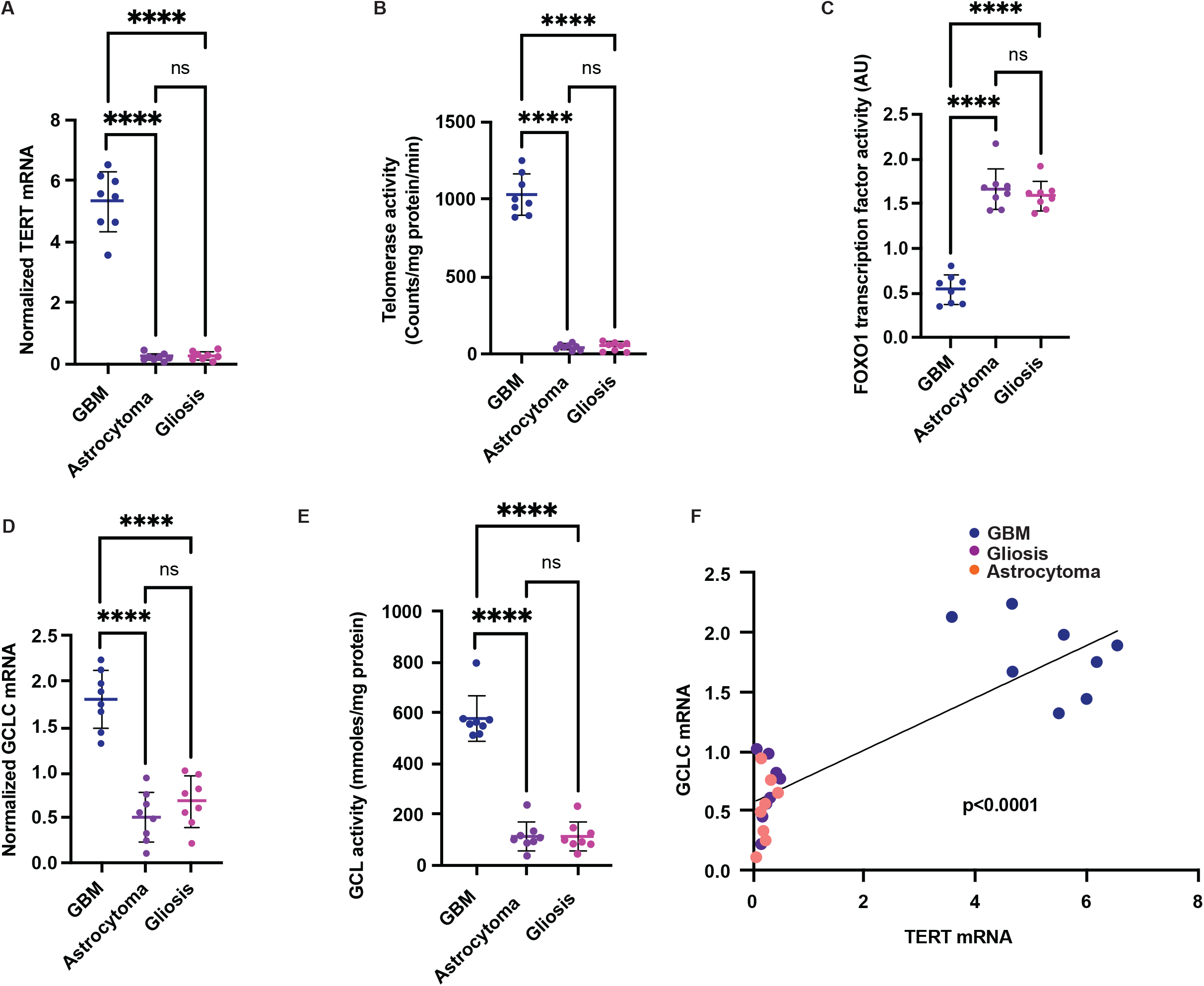
TERT expression is associated with elevated GCLC expression and GCL activity in patient biopsies. TERT mRNA **(A)**, telomerase activity **(B)**, FOXO1 transcription factor activity **(C)**, GCLC mRNA **(D)**, and GCL activity **(E)** in GBM, astrocytoma, or gliosis biopsies. **(F)** Linear correlation between TERT mRNA and GCLC mRNA in GBM, astrocytoma, or gliosis biopsies. ** represents p<0.01, *** represents p<0.001 and **** represents p<0.0001.

### GCLC inhibition rewires [U-^13^C]-glutamine metabolism and modestly reduces the clonogenicity of GBM cells

Next, we examined the therapeutic potential of targeting GCLC in GBM cells. To this end, we used siRNA to silence GCLC in both GBM6 and U251 models (" ref-type="fig">Fig. 3A-3B). As shown in " ref-type="fig">Fig. 3C-3E, GCLC silencing increased levels of ROS and modestly reduced clonogenicity but did not induce apoptosis. We also examined the effect of pharmacologically inhibiting GCLC using buthionine sulfoximine (BSO) (30). BSO significantly reduced GCL activity, and inhibited clonogenicity with an IC50 of 7.84 μM for U251 and 12.64 μM for GBM6 cells (" ref-type="fig">Fig. 3F-3G). As with siRNA-mediated GCLC silencing, although BSO increased ROS generation, it did not induce apoptosis (" ref-type="fig">Fig. 3H-3I).

**Figure 3.**
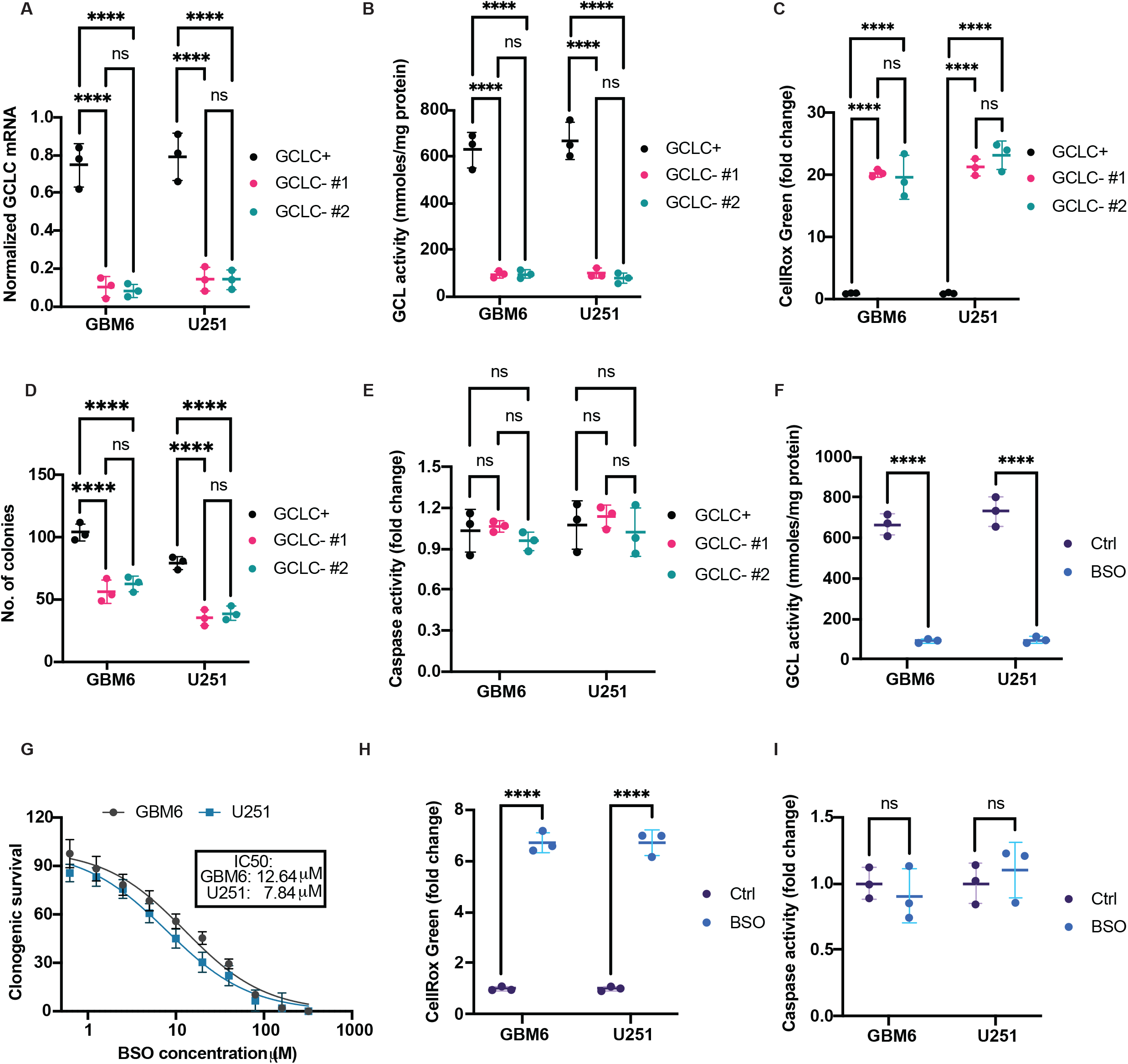
GCLC silencing or inhibition induces oxidative stress and modestly inhibits clonogenicity in GBM cells. Verification of reduced GCLC mRNA **(A)** and GCL activity **(B)** in GBM6 and U251 cells transfected with non-overlapping siRNA against GCLC. Effect of GCLC silencing on levels of ROS **(C)**, soft agar colony formation **(D)**, and caspase activity **(E)** in GBM6 and U251 cells. **(F)** Verification of reduced GCL activity in GBM6 and U251 cells treated with vehicle (DMSO) or 10 μM BSO for 48 h. **(G)** Dose response curve of BSO for GBM6 and U251 cells. Cells were treated with the indicated concentrations of BSO and IC50 measured via % inhibition of clonogenicity. Effect of BSO on levels of ROS **(H)** and caspase activity **(I)** in GBM6 and U251 cells. ** represents p<0.01, *** represents p<0.001 and **** represents p<0.0001.

To assess the metabolic consequences of GCLC inhibition and identify potential compensatory metabolic alterations, we examined [U-^13^C]-glutamine metabolism (see schematic in Fig. 4A) in our models. In addition to GSH synthesis, glutamine-derived glutamate can be transaminated to α-ketoglutarate (α-KG), which can then be metabolized via the TCA cycle to produce oxaloacetate, and aspartate (31,38,39). Glutamine, aspartate, and bicarbonate are converted by a single trifunctional, rate-limiting enzyme called carbamoyl phosphate synthetase 2, aspartate transcarbamylase, and dihydroorotase (CAD) to dihydroorotate (40). Subsequent metabolism generates the pyrimidine nucleotides uridine triphosphate (UTP) and cytidine triphosphate (CTP). Alternately, glutamine can shunted towards purine biosynthesis and be incorporated into guanosine monophosphate (GMP) and adenosine monophosphate (AMP) (40). As shown in " ref-type="fig">Fig. 4B-4E, GCLC silencing and BSO reduced GSH synthesis from [U-^13^C]-glutamine, confirming on-target activity in both GBM6 and U251 models. However, GCLC inhibition upregulated ^13^C labeling of glutamate and oxidative metabolism of [U-^13^C]-glutamine via the TCA cycle to m+4 α-KG, succinate, and malate (" ref-type="fig">Fig. 4A-4D). We did not observe any change in m+5 citrate, indicating that reductive glutamine metabolism was not altered by GCLC inhibition (39,41). Importantly, GCLC inhibition increased m+4 aspartate, m+4 dihydroorotate, and m+3 UTP and CTP (" ref-type="fig">Fig. 4A-4D). We confirmed that these differences in dynamic glutamine metabolism resulted in corresponding differences in metabolite pool sizes i.e., reduced GSH abundance, and elevated glutamate, TCA cycle, aspartate, and pyrimidine nucleotides (Supplementary Fig. S2A-S2D). GCLC inhibition did not impact purine nucleotide abundance or synthesis from [U-^13^C]-glutamine in either model (see " ref-type="fig">Fig. 4B-4E; Supplementary Fig. S2A-S2D). Taken together, these results suggest that GCLC inhibition reduces GSH synthesis while upregulating oxidative glutamine metabolism and pyrimidine nucleotide biosynthesis in GBM cells.

**Figure 4.**
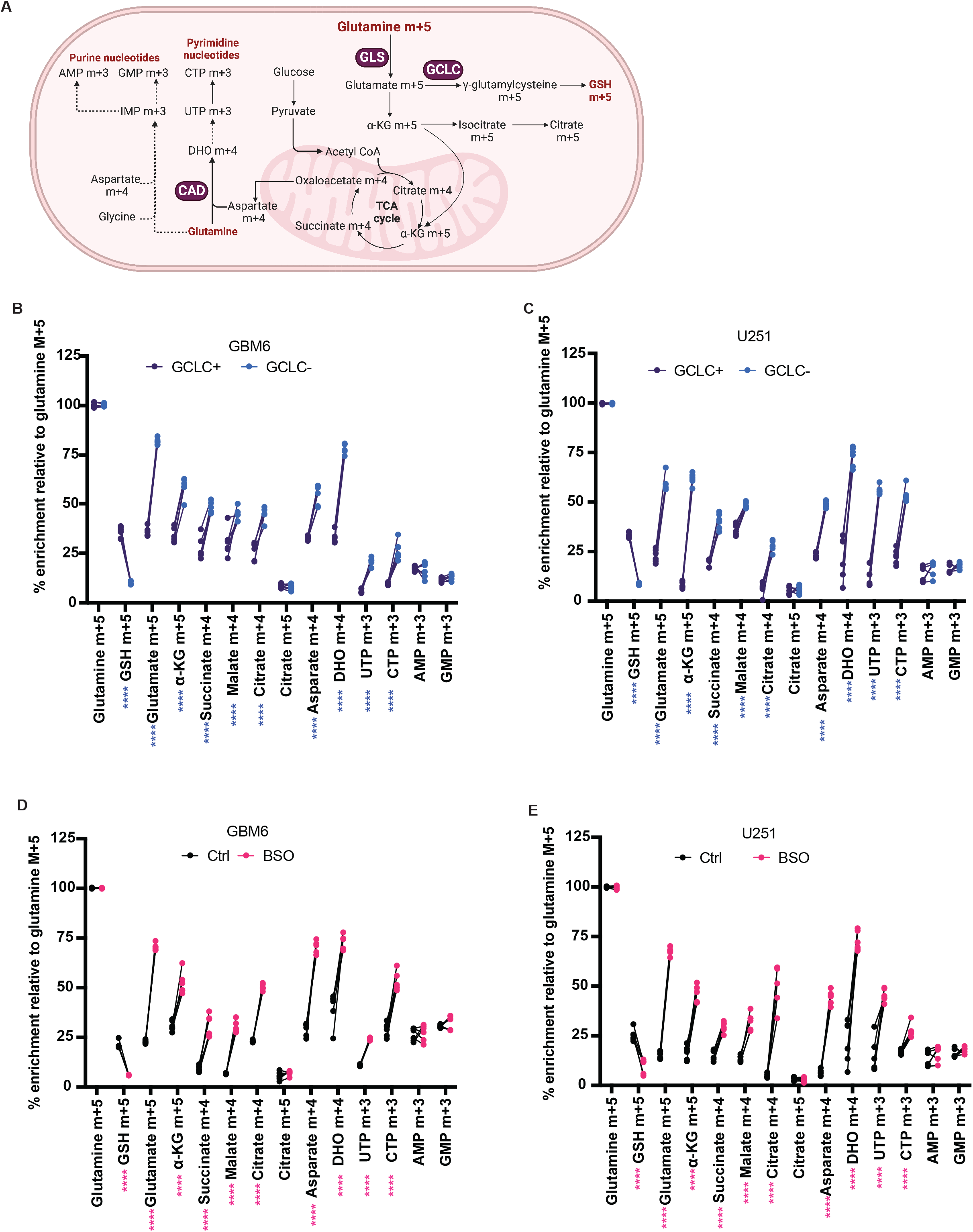
GCLC silencing or inhibition rewires [U-^13^C]-glutamine metabolism in GBM cells. (**A**) Schematic illustration of [U-^13^C]-glutamine metabolism in cancer cells. [U-^13^C]-glutamine (m+5 glutamine) is catabolized by GLS to m+5 glutamate, which can then be converted by GCLC to m+5 ℽ-glutamyl cysteine and then to m+5 GSH. Glutamate m+5 can also be transaminated to m+5 α-KG by transaminases such as *GLUD1*, *GLUD2*, *BCAT1*, and *GPT1*. α-KG m+5 is oxidized via the TCA cycle to m+4 succinate, m+4 malate, and m+4 oxaloacetate. Oxaloacetate m+4 is conjugated with acetyl CoA derived from glucose to form m+4 citrate. In addition to oxidative metabolism, m+5 α-KG can be reductively carboxylated by IDH1 to m+5 isocitrate and then m+5 citrate. Alternately, m+4 oxaloacetate is converted by *GOT1* to m+4 aspartate. CAD combines aspartate, glutamine, and bicarbonate to produce m+4 dihydroorotate (DHO), which is subsequently metabolized by *DHODH* and *UMPS* to the pyrimidine nucleotides m+3 UTP and m+3 CTP. Aspartate m+4 along with glutamine and glycine also produces the purine nucleotides m+3 AMP and m+3 GMP. Effect of GCLC silencing using siRNA on [U-^13^C]-glutamine metabolism in GBM6 **(B)** or U251 **(C)** cells. Effect of inhibiting GCLC using BSO on [U-^13^C]-glutamine metabolism in GBM6 **(D)** or U251 **(E)** cells. ** represents p<0.01, *** represents p<0.001 and **** represents p<0.0001.

### GCLC inhibition upregulates GLS and CAD in a MYC-dependent manner in GBMs

Next, we examined the mechanism by which GCLC inhibition rewires glutamine metabolism in GBMs. Expression profiling of enzymes involved in glutamine metabolism showed that GCLC inhibition significantly upregulated expression of GLS and CAD (" ref-type="fig">Fig. 5A-5D), which are the rate-limiting enzymes for glutamate and dihydroorotate synthesis (see schematic illustration in Fig. 4A) (38,40). MYC is a known transcriptional regulator of both GLS and CAD (42,43). As shown in " ref-type="fig">Fig. 5E-5F, both GCLC silencing and BSO treatment upregulated MYC expression in the GBM6 and U251 models. Importantly, silencing MYC using siRNA in GCLC– or BSO-treated cells (Supplementary Figure S3A-S3B) downregulated expression of GLS and CAD to levels observed in GCLC+ cells (" ref-type="fig">Fig. 5G-5J). Collectively, these results suggest that GCLC inhibition upregulates MYC, which drives expression of GLS and CAD in GBM cells.

**Figure 5.**
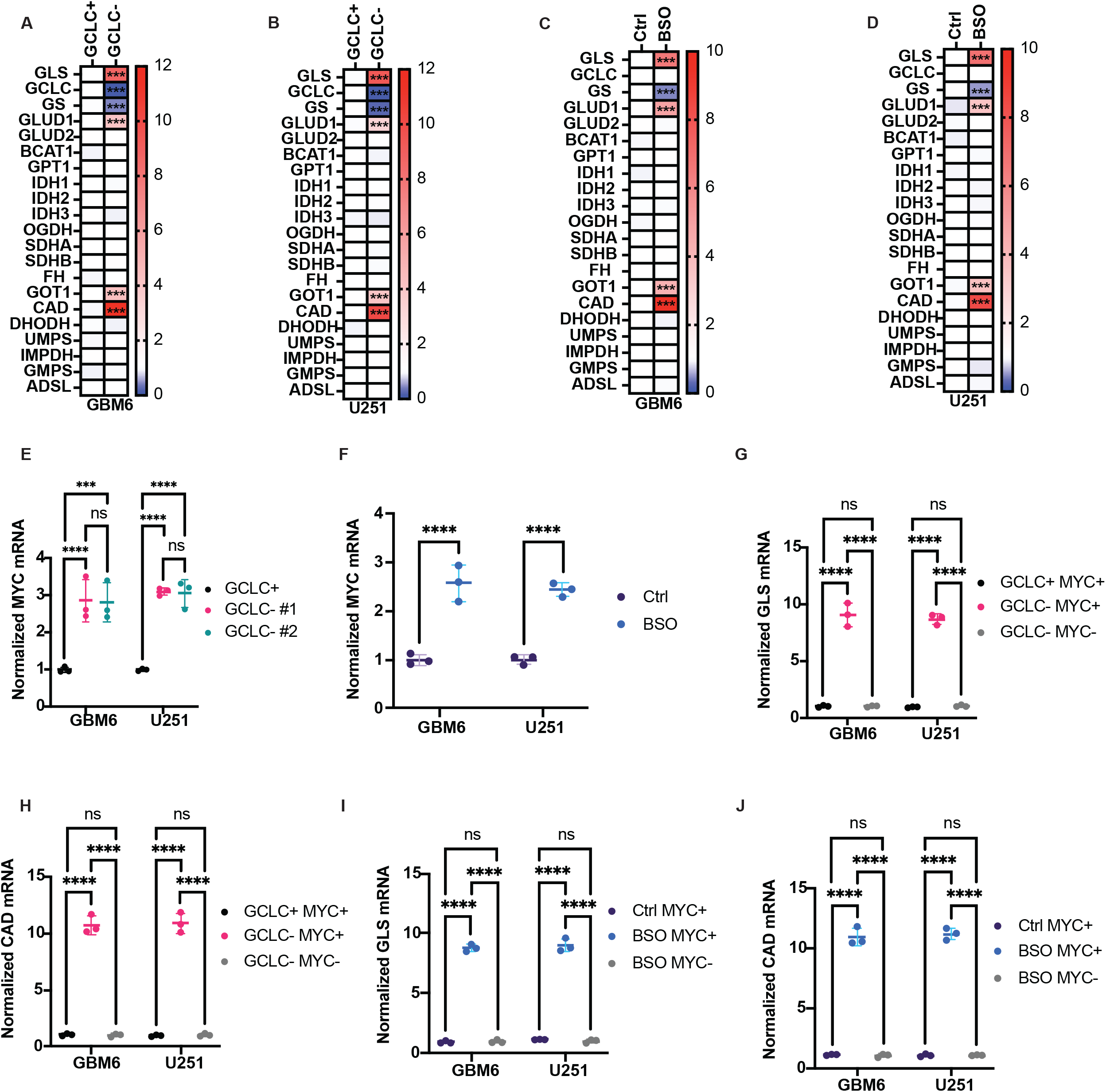
GCLC silencing or inhibition upregulates GLS and CAD in a MYC-dependent manner in GBM cells. Effect of GCLC silencing by siRNA on mRNA levels of enzymes involved in glutamine metabolism in GBM6 **(A)** and U251 **(B)** cells. Effect of GCLC inhibition using BSO on mRNA levels of enzymes involved in glutamine metabolism in GBM6 **(C)** and U251 **(D)** cells. Effect of GCLC silencing by siRNA **(E)** or GCLC inhibition by BSO **(F)** on expression of MYC in GBM6 and U251 cells. mRNA levels of GLS **(G)** and CAD **(H)** in GCLC+ and GCLC-GBM6 and U251 cells. To probe for the role of MYC in upregulating GLS and CAD following GCLC silencing, we examined GCLC-cells in which MYC was silenced by siRNA. mRNA levels of GLS **(I)** and CAD **(J)** in GBM6 and U251 cells treated with vehicle (DMSO) or 10 μM BSO for 48 h. To probe for the role of MYC in upregulating GLS and CAD following GCLC inhibition, we examined silenced MYC by siRNA concurrently with BSO treatment. ** represents p<0.01, *** represents p<0.001 and **** represents p<0.0001.

### Inhibiting GLS and CAD is synthetically lethal in combination with GCLC inhibition in GBM cells

Since our data indicated that GCLC inhibition upregulated expression of GLS and CAD, we examined the therapeutic potential of targeting GLS and CAD in combination with GCLC in GBM cells. 6-diazo-5-oxy-L-norleucin (DON) is a potent inhibitor of glutamine-utilizing enzymes including GLS and the carbamoyl phosphate synthetase II domain of CAD (44). First, we confirmed that DON as monotherapy significantly downregulated GLS and CAD activity and inhibited clonogenicity with an IC50 of 6.27 μM for GBM6 and 5.24 μM for U251 cells (" ref-type="fig">Fig. 6A-6C). Examination of [U-^13^C]-glutamine metabolism showed that DON significantly downregulated oxidative metabolism to m+5 glutamate, m+5 α-KG, m+4 succinate, m+4 malate, and m+4 aspartate, consistent with inhibition of GLS activity (" ref-type="fig">Fig. 6D-6E). DON also downregulated ^13^C labeling of m+4 dihydroorotate, m+3 UTP and m+3 CTP, as expected with inhibition of CAD activity, in both GBM6 and U251 models (" ref-type="fig">Fig. 6D-6E). DON did not alter reductive metabolism of [U-^13^C]-glutamine to m+5 citrate or to m+3 AMP or GMP (" ref-type="fig">Fig. 6D-6E). DON reduced steady-state pool sizes of TCA cycle metabolites and pyrimidine nucleotides in both GBM6 and U251 models (" ref-type="fig">Fig. 6F-6G). However, despite on-target activity, DON as monotherapy did not induce cell death in either model (Supplementary Fig. S3C).

**Figure 6.**
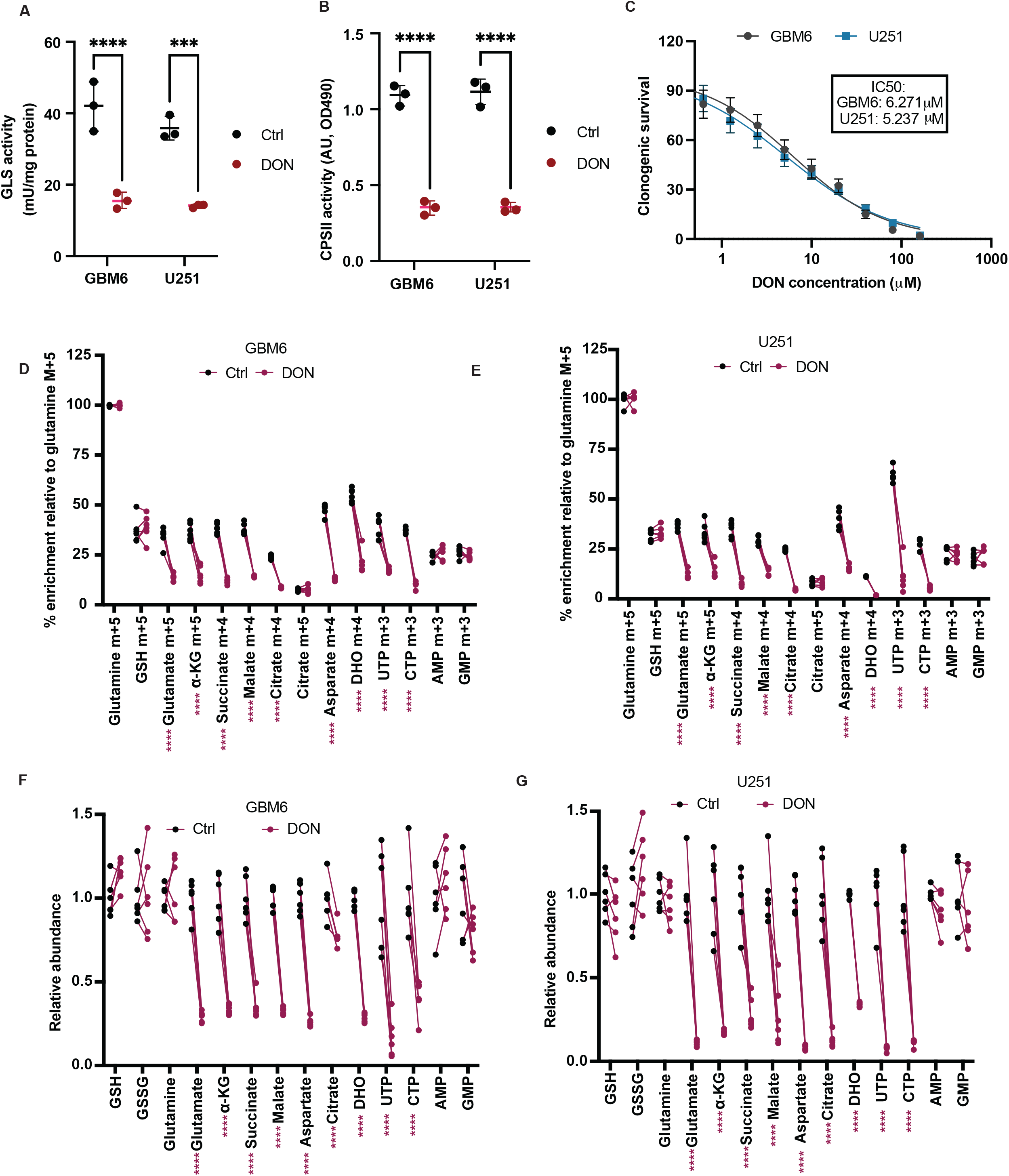
DON inhibits GLS and CAD and downregulates pyrimidine nucleotide biosynthesis from [U-^13^C]-glutamine in GBM cells. Effect of DON on GLS activity **(A)** and CAD activity **(B)** in GBM6 and U251 cells. **(C)** Dose response curve of DON for GBM6 and U251 cells. Cells were treated with the indicated concentrations of DON and IC50 measured via % inhibition of clonogenicity. Effect of DON on % ^13^C metabolite enrichment from [U-^13^C]-glutamine in GBM6 **(D)** or U251 **(E)** cells. Effect of DON on metabolite pool sizes in GBM6 **(F)** or U251 **(G)** cells. ** represents p<0.01, *** represents p<0.001 and **** represents p<0.0001.

Next, we examined the effect of combined treatment with BSO and DON in GBM6 and U251 cells. As shown in " ref-type="fig">Fig. 7A-7B, the combination of BSO and DON was synergistically lethal with a Bliss synergy score of 31.37 for the GBM6 model and 29.19 for the U251 model (>10 indicates synergy). With both models, DON induced a synergistic potency shift of IC50 for BSO, and BSO induced a synergistic potency shift of IC50 for DON (Supplementary Fig. S4A-S4B). We next examined the effect of treatment with a combination of 1 μM each of DON and BSO (based on the dose-response matrix of the combination; see Supplementary Fig. S4C-S4D). As shown in Fig. 7C, the combination of DON and BSO induced apoptosis in both GBM6 and U251 cells. To confirm the mechanism of action, we examined [U-^13^C]-glutamine metabolism in GBM6 and U251 cells treated with vehicle or the combination of DON and BSO. As shown in " ref-type="fig">Fig. 7D-7E, the combination of DON and BSO abrogated synthesis of m+5 GSH, m+5 glutamate, m+5 α-KG, m+4 succinate, m+4 malate, m+4 aspartate, m+4 dihydroorotate, m+3 UTP and m+3 CTP. Concomitantly, pool sizes of these metabolites were also significantly reduced in GBM6 and U251 cells treated with the combination of BSO and DON relative to vehicle (" ref-type="fig">Fig. 7F-7G). Taken together, these results highlight the synthetic lethality of treatment with a combination of BSO and DON in GBM cells.

**Figure 7.**
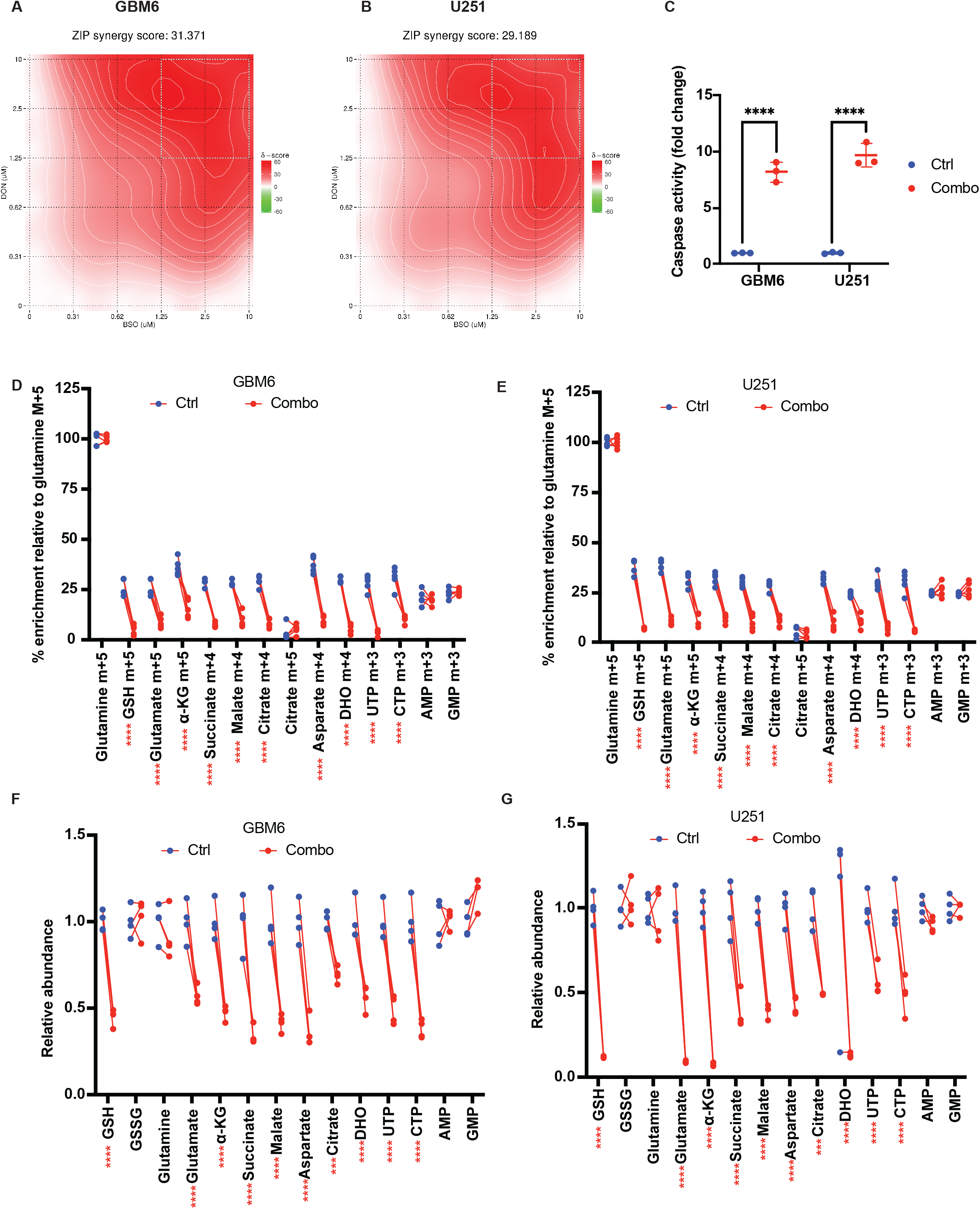
Combined inhibition of GCLC, GLS and CAD is synthetically lethal in GBM cells. Bliss synergy score maps for the combination of DON and BSO in GBM6 **(A)** and U251 **(B)** cells. **(C)** Effect of treatment with vehicle (Ctrl) or the combination of 1 μM each of DON and BSO (Combo) on caspase activity in GBM6 and U251 cells. Effect of treatment with vehicle (Ctrl) or the combination of 1 μM each of DON and BSO (Combo) on % ^13^C metabolite enrichment from [U-^13^C]-glutamine in GBM6 **(D)** or U251 **(E)** cells. Effect of treatment with vehicle (Ctrl) or the combination of 1 μM each of DON and BSO (Combo) on metabolite pool sizes in GBM6 **(F)** or U251 **(G)** cells. ** represents p<0.01, *** represents p<0.001 and **** represents p<0.0001.

### Combined inhibition of GCLC, GLS and CAD induces tumor shrinkage in preclinical GBM models *in vivo*

The use of DON has been associated with disease stabilization or remission in clinical trials in cancer patients (44). However, clinical use was halted due to issues of gastrointestinal toxicity (44). JHU-083 is a novel prodrug of DON that has shown efficacy in multiple cancers, including a cytostatic effect in preclinical GBM models (29,39). JHU-083 is also brain penetrant, which is an important consideration for GBM therapy (29). We first confirmed that JHU-083 inhibits GLS and CAD activity in both GBM6 and U251 cells (Supplementary Fig. S5A-S5B). Like DON, JHU-083 as monotherapy did not induce apoptosis (Supplementary Fig. S5C). We then examined the effect of treatment with vehicle or the combination of JHU-083 and BSO on mice bearing intracranial patient-derived GBM6 tumors. As shown in " ref-type="fig">Fig. 8A-8B, the combination of JHU-083 and BSO induced tumor shrinkage and significantly extended survival. Importantly, *in vivo* infusion with [U-^13^C]-glutamine showed a significant reduction in synthesis and steady-state levels of GSH, glutamate, α-KG, succinate, malate, aspartate, dihydroorotate, UTP and CTP in GBM6 tumor-bearing mice treated with the combination of JHU-083 and BSO relative to controls (" ref-type="fig">Fig. 8C-8D). Concomitantly, GCL, GLS and CAD activity were significantly downregulated in tumor tissue from GBM6 tumor-bearing mice treated with the combination of JHU-083 and BSO relative to controls, further confirming on-target activity (" ref-type="fig">Fig. 8E-8G). These results indicate that combined inhibition of GCLC, GLS and CAD is synthetically lethal *in vitro* and *in vivo,* and highlight BSO and JHU-083 as potential therapeutic agents for GBMs.

**Figure 8.**
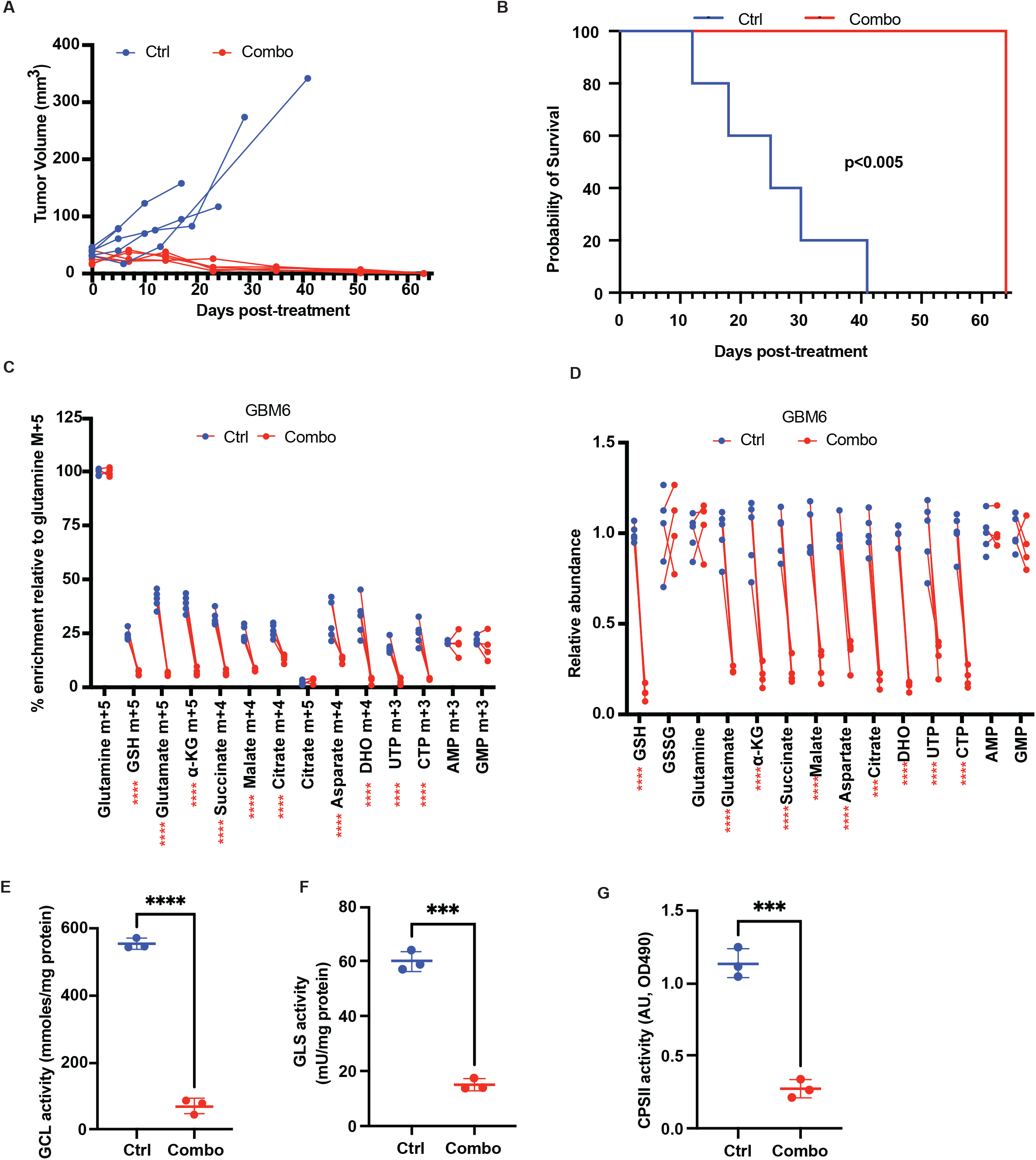
Combined inhibition of GCLC, GLS and CAD causes tumor shrinkage in mice bearing intracranial GBM xenografts *in vivo*. Effect of treatment with vehicle or the combination of 20 mg/kg each of JHU-083 and BSO on tumor volume over time **(A)** and animal survival **(B)** in mice bearing intracranial GBM6 tumors. % ^13^C metabolite enrichment from [U-^13^C]-glutamine **(C)** and metabolite pool sizes **(D)** in tumor tissue from mice bearing intracranial GBM6 tumors treated with vehicle or the combination of 20 mg/kg each of JHU-083 and BSO. Mice were treated for 7 days, infused with [U-^13^C]-glutamine, and tumor tissue resected for LC-MS. GCL activity **(E),** GLS activity **(F),** and CAD activity **(G)** in tumor tissue from mice bearing intracranial GBM6 tumors treated with vehicle or the combination of 20 mg/kg each of JHU-083 and BSO. ** represents p<0.01, *** represents p<0.001 and **** represents p<0.0001.

## DISCUSSION

TERT expression is essential for tumor initiation and maintenance in ∼85% of all human cancers (4,5). It is silenced early in development in normal somatic cells, with the exception of stem cells, and reactivated specifically in tumor cells via hotspot mutations in the TERT promoter (5,7). Studies suggest that TERT promoter mutations occur years before initial diagnosis (45,46) and that TERT expression is potentially clonal in GBMs (47). While these criteria make TERT an attractive therapeutic target, targeting TERT directly has been fraught with difficulty due to issues of stem cell toxicity (4,6,7). Furthermore, the lag period before telomere shortening occurs following TERT inhibition is problematic for aggressively proliferating tumors such as GBMs (4,6,7). Dissecting the metabolic reprogramming that is associated with TERT expression provides targets that are potentially druggable. Our studies show that TERT drives GSH synthesis by upregulating GCLC in GBM cells. However, targeting GCLC upregulates GLS and CAD, the rate-limiting enzymes for glutamine metabolism via the TCA cycle and pyrimidine nucleotide biosynthesis. Importantly, combined inhibition of GCLC, GLS and CAD is synergistic and induces tumor regression in mice bearing intracranial GBMs *in vivo*.

Replicative immortality and metabolic reprogramming are both hallmarks of cancer (48). TERT expression drives replicative immortality in cancer (4,5). Tumors also rewire metabolism to maintain redox balance and mitigate oxidative stress (49). Our studies provide a mechanistic framework that integrates TERT expression with reprogramming of GSH metabolism in GBMs. By combining loss-of-function experiments with metabolomics and stable isotope tracing in cells and *in vivo* in mice bearing intracranial tumor xenografts, we provide direct evidence linking TERT expression with elevated GSH synthesis from [U-^13^C]-glutamine in GBMs. Our studies identify a mechanistic role for the transcription factor FOXO1 in TERT-mediated upregulation of GCLC, the catalytic component of GCL, which is the rate-limiting enzyme for GSH synthesis. Targeting GCLC using siRNA or BSO downregulates GSH synthesis and inhibits clonogenicity. However, GCLC inhibition upregulates pyrimidine nucleotide biosynthesis from [U-^13^C]-glutamine, an effect that is driven by upregulation of GLS and CAD in a MYC-dependent manner. Importantly, combined inhibition of GCLC, GLS, and CAD is synthetically lethal *in vitro* and *in vivo.* These findings highlight the plasticity that is built into metabolic networks and point to the utility of *in vivo* stable isotope tracing in unraveling the mechanisms by which tumors bypass GCLC inhibition.

Our studies underscore metabolic synthetic lethality as a promising approach for cancer therapy. We find that the combination of BSO and JHU-083 causes tumor shrinkage in mice bearing patient-derived GBM xenografts *in vivo.* Regarding clinical translation, BSO is limited by poor brain penetrance and pharmacokinetic properties (50). Our studies provide a rational impetus for the development of novel GCLC inhibitors with improved pharmacokinetic properties. JHU-083 is a pro-drug of DON that was designed to cross the blood brain barrier and get selectively activated within the tumor, leading to inhibition of glutamine-utilizing enzymes, including GLS and CAD (29,39,44). Of note, a related DON pro-drug (DRP-104) that is not brain penetrant is in clinical trials for solid tumors (NCT06027086) (51). Our results indicate that both DON and JHU-083 inhibit GLS and CAD activity in GBM cells. Consistent with these results, treatment with DON as monotherapy reduced ^13^C labeling of TCA cycle metabolites and the pyrimidine nucleotides UTP and CTP. Interestingly, despite reports suggesting that DON inhibits the purine biosynthesis enzymes formylglycinamide ribonucleotide amidotransferase and GMP synthetase (44), we did not observe inhibition of purine nucleotide (AMP and GMP) biosynthesis following treatment with DON or JHU-083 in our models. It is possible that differences in the affinity of DON/JHU-083 to different glutamine-utilizing enzymes result in context-dependent differences in the metabolic consequences of DON/JHU-083 treatment. Nevertheless, our studies emphasize the therapeutic potential of combination therapies targeting GCLC, GLS and CAD for the treatment of GBMs.

In summary, our studies leverage a mechanistic understanding of TERT-associated metabolic rewiring for the identification of synthetically lethal metabolic vulnerabilities in GBMs. Clinical translation of our results has the potential to enable the rational development of novel combination therapies for GBM patients.

## AUTHOR CONTRIBUTIONS

SU, CT, MT, GB, AMG, and PV performed and analyzed the experiments; SMR contributed to the acquisition of the LC-MS system; JtH and TGG provided technical and intellectual support for the metabolomics and stable isotope tracing studies; PV conceptualized the research, directed the studies, and secured funding.

---

**Supplementary Figure 1.**
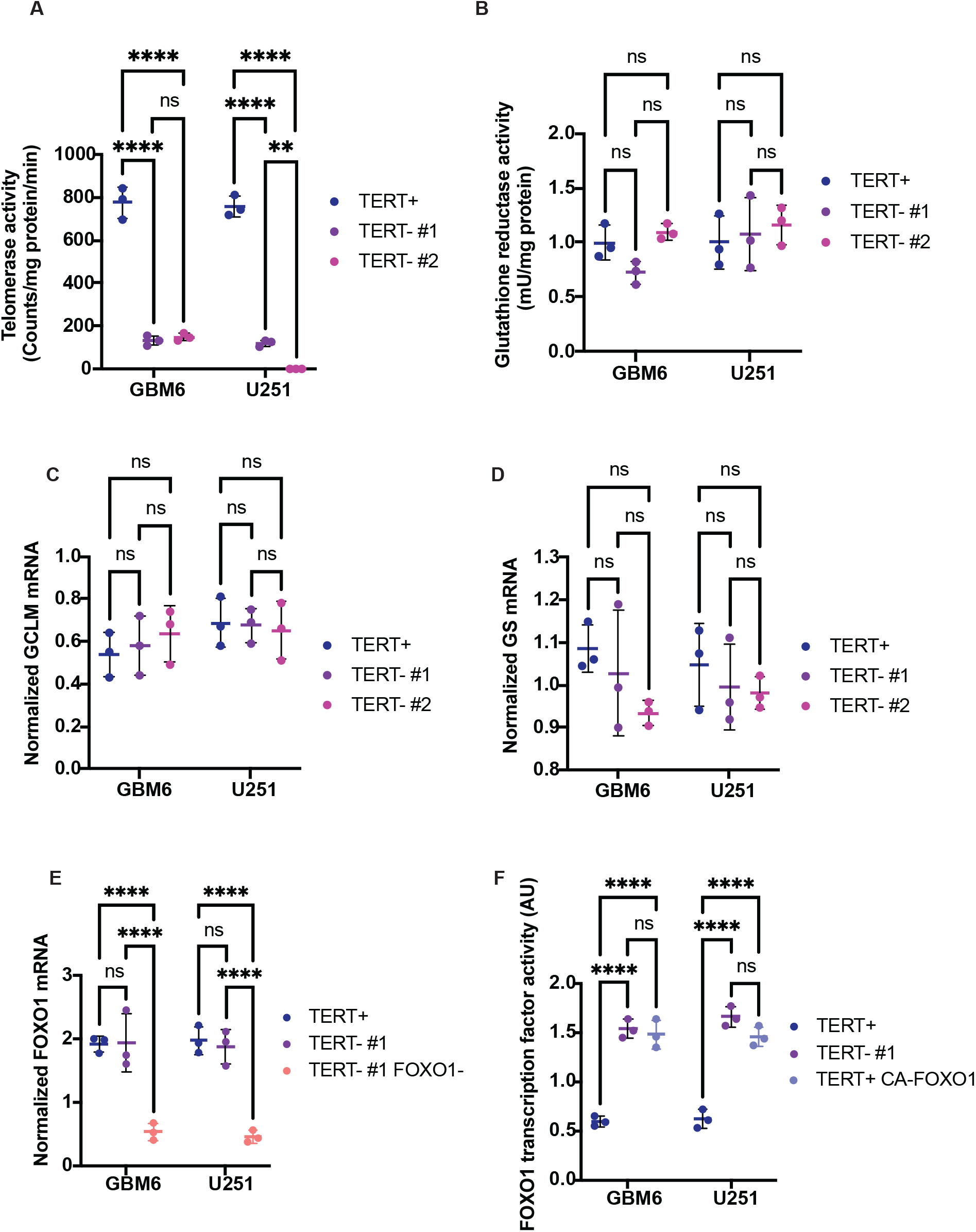
Verification of gene silencing in GBM cells. Telomerase activity **(A)**, glutathione reductase activity **(B)**, GCLM mRNA **(C)**, and glutathione synthetase mRNA **(D)** in GBM6 and U251 cells transfected with non-targeting siRNA (TERT+) or siRNA against TERT (TERT-). **(E)** Verification of loss of FOXO1 mRNA in GBM6 and U251 cells transfected with non-targeting siRNA (TERT+), siRNA against TERT (TERT-#1), or siRNA against both TERT and FOXO1 (TERT-+1 FOXO1-). **(F)** Verification of FOXO1 transcription factor activity in GBM6 and U251 cells transfected with non-targeting siRNA (TERT+), siRNA against TERT (TERT-#1), or with a plasmid expressing a constitutively active form of FOXO1 (TERT+ CA-FOXO1). ** represents p<0.01, *** represents p<0.001, **** represents p<0.0001 and ns represents non-significance.

**Supplementary Figure 2.**
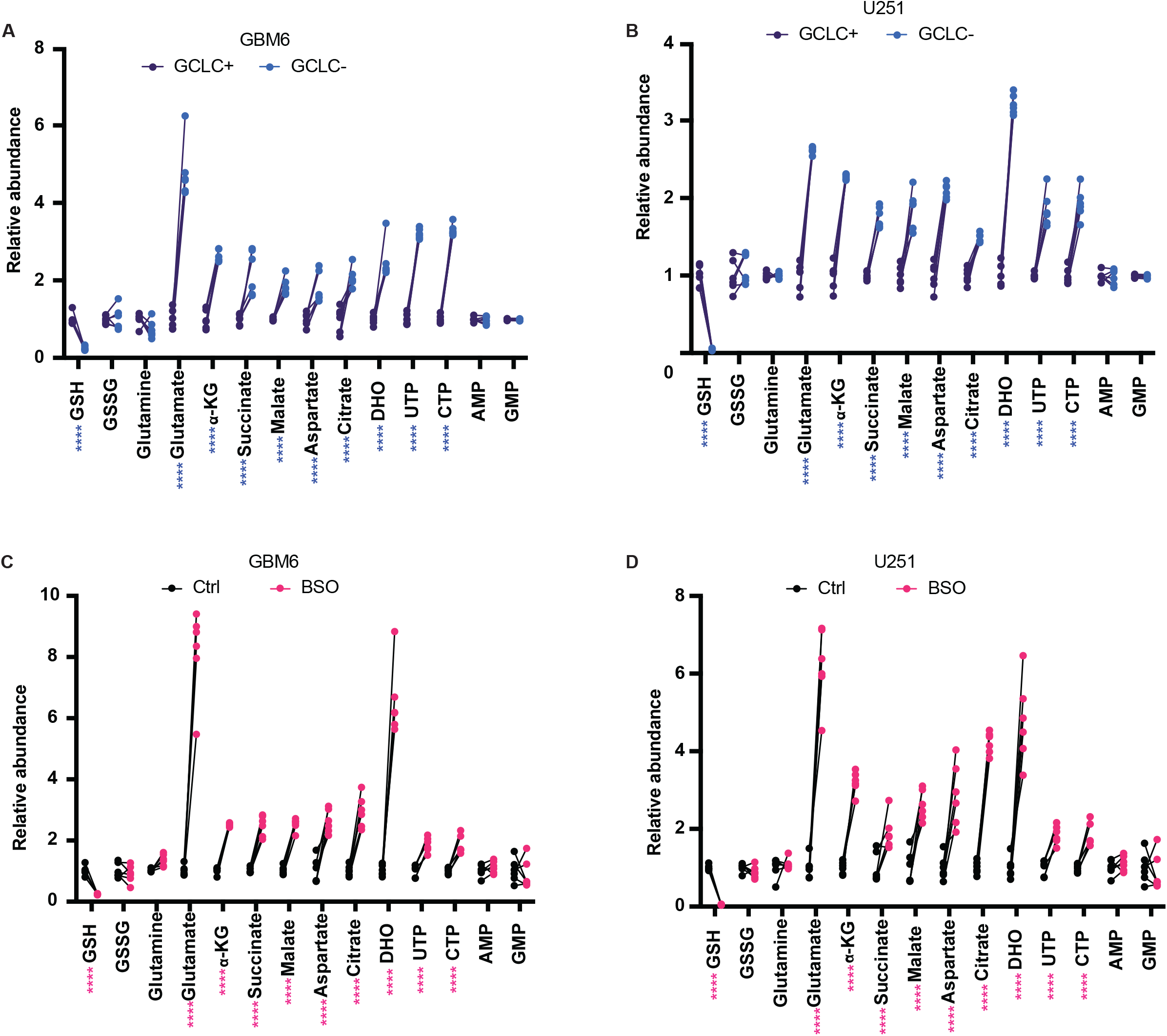
GCLC inhibition reduces GSH pool size but upregulates levels of TCA cycle metabolites and pyrimidine nucleotides. Effect of silencing GCLC using siRNA on steady-state metabolite pool sizes in the GBM6 **(A)** and U251 **(B)** models. Effect of inhibiting GCLC using BSO on steady-state metabolite pool sizes in the GBM6 **(C)** and U251 **(D)** models. ** represents p<0.01, *** represents p<0.001, **** represents p<0.0001 and ns represents non-significance.

**Supplementary Figure 3.**
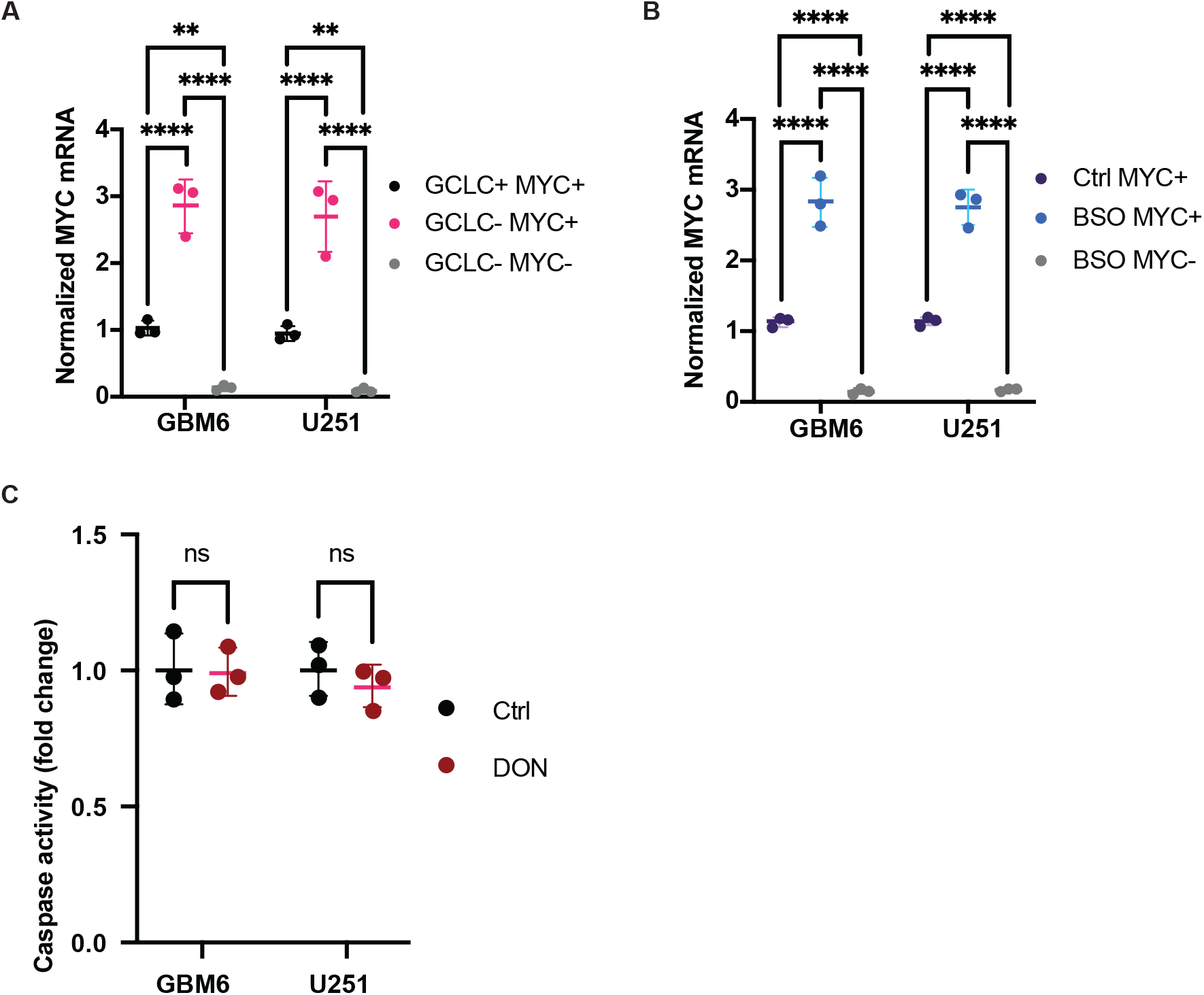
Verification of gene silencing in GBM cells. (**A**) Verification of loss of MYC mRNA in GBM6 and U251 cells transfected with non-targeting siRNA (GCLC+ MYC+), siRNA against GCLC (GCLC-MYC+), or siRNA against both GCLC and MYC (GCLC-MYC-). **(B)** Verification of loss of MYC mRNA in GBM6 and U251 cells treated with vehicle (DMSO), 10 μM BSO (BSO MYC+) or 10 μM BSO concurrently with siRNA against MYC (BSO MYC-). **(C)** Effect of 5 μM DON on caspase activity in GBM6 and U251 cells. ** represents p<0.01, *** represents p<0.001, **** represents p<0.0001 and ns represents non-significance.

**Supplementary Figure 4.**
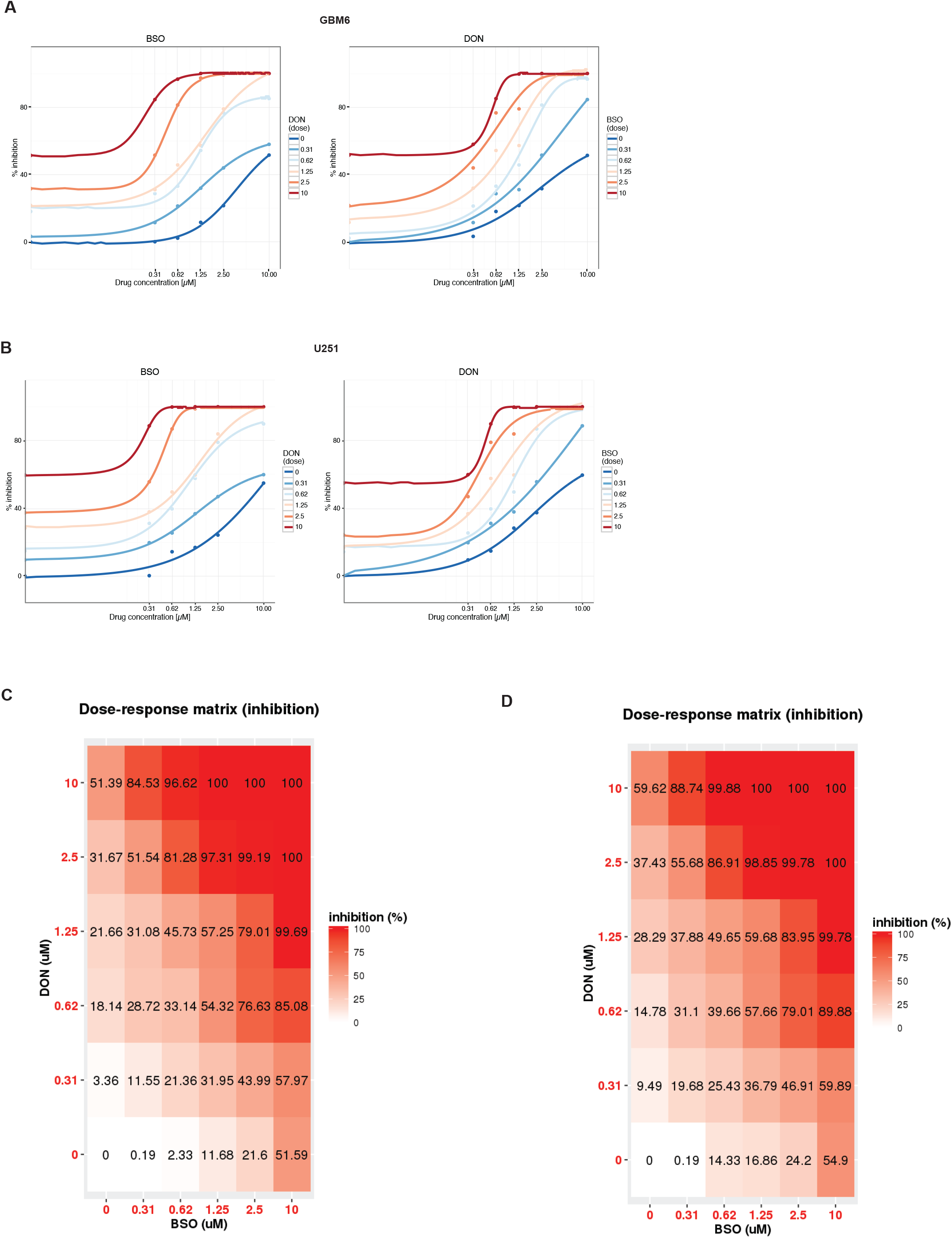
Combined inhibition of GCLC, GLS and CAD is synthetically lethal in GBM cells. **(A)** Dose response curves showing the shift in potency (IC50) caused by addition of the indicated concentrations of DON to GBM6 cells treated with BSO (left panel) or by the addition of the indicated concentrations of BSO to GBM6 cells treated with DON (right panel). **(B)** Dose response curves showing the shift in potency (IC50) caused by addition of the indicated concentrations of DON to U251 cells treated with BSO (left panel) or by the addition of the indicated concentrations of BSO to U251 cells treated with DON (right panel). **(C)** Dose response matrix of % inhibition of clonogenicity caused by treatment with a serial dilution of DON and BSO at the indicated concentrations for the GBM6 model. **(D)** Dose response matrix of % inhibition of clonogenicity caused by treatment with a serial dilution of DON and BSO at the indicated concentrations for the U251 model.

**Supplementary Figure 5.**
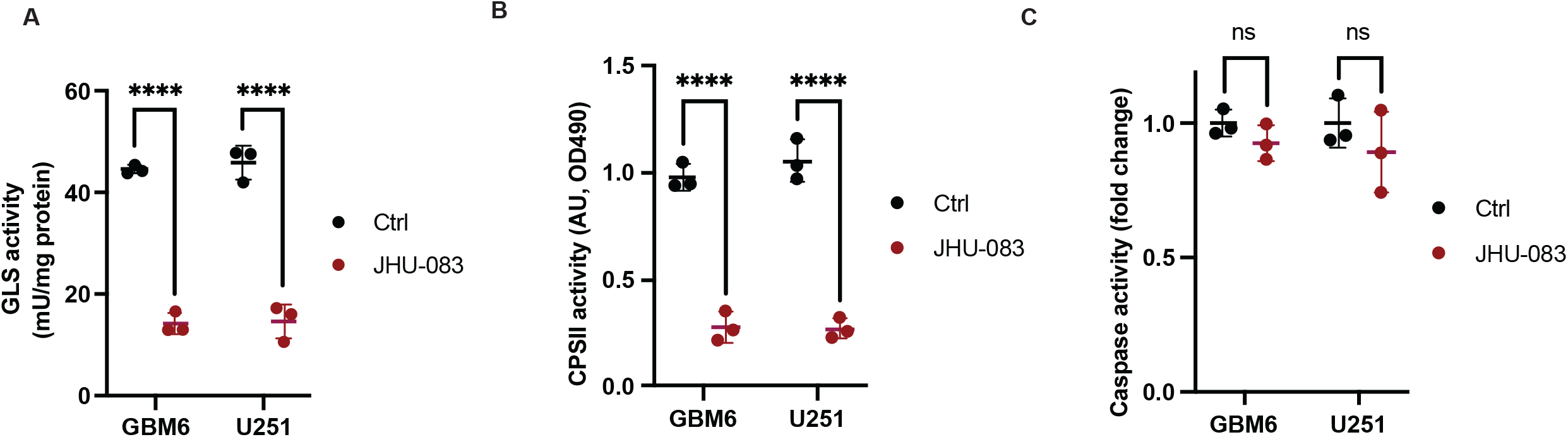
JHU-083 inhibits GLS and CAD in GBM cells. Effect of treatment with vehicle (DMSO) or 1 μM JHU-083 on GLS activity **(A)**, CAD activity **(B)**, and caspase activity **(C)** in GBM6 and U251 cells.

## Notes

### Competing Interest Statement

TGG has consulting and equity agreements with Auron Therapeutics, Boundless Bio, Coherus BioSciences and Trethera Corporation. The other authors have no conflicts of interest to declare.

